# A multivirulent *Plasmopara viticola* strain from Cilaos on Réunion Island breaks down *Rpv1, Rpv3.1 and Rpv10* mediated resistance of grapevine

**DOI:** 10.1101/2025.09.02.673614

**Authors:** Julie Ramírez Martínez, Anne-Sophie Miclot, Etienne Dvorak, Isabelle D. Mazet, Carole Couture, Laurent Delière, Frédéric Fabre, Ignace Hoarau, Olivier Yobrégat, Marie Foulongne-Oriol, François Delmotte

## Abstract

Grapevine downy mildew, caused by *Plasmopara viticola*, is one of the most destructive diseases in viticulture. Resistance-based management strategies rely on grapevine varieties carrying major resistance loci (*Rpv*). Although breakdown of several loci has been reported, *Rpv1* had remained effective until now. Here, we provide the first evidence of *Rp*v1 breakdown by *P. viticola* in Cilaos on Réunion Island (France) and the first case of a strain simultaneously overcoming three resistances of grapevine. We combined pathogenicity assays with whole-genome sequencing to characterize a *P. viticola* strain collected in Cilaos in 2023, alongside a panel of eight reference strains of known virulence. Hypersensitive response and sporulation were assessed for each strain on Chardonnay (susceptible variety) and four resistant varieties carrying *Rpv1*, *Rpv3.1*, *Rpv10,* or *Rpv12*. The strain collected in Cilaos was able to overcome not only *Rpv1* but also *Rpv3.1* and *Rpv10*. On *Rpv1*, we observed a complete loss of host recognition with high sporulation. On *Rpv3.1*, the phenotype was consistent with previous breakdowns, and the strain carried the *vir1* allele previously described in France. By contrast, the breakdown of *Rpv10* differed from that reported in European populations: whereas European strains displayed only partial breakdown of resistance, the strain of Cilaos showed complete loss of host recognition with high sporulation. Moreover, genomic analyses revealed a novel mutation, a large homozygous deletion in the corresponding *avr* locus. Our genomic data analyses further suggests that this *P. viticola* strain shares a genetic background with populations from mainland France, raising serious concerns about the potential emergence and spread of multivirulent lineages in Europe. These findings highlight the need for large-scale virulence monitoring of *P. viticola* and improved strategies for the sustainable management of grapevine resistance in Europe.

## Introduction

Grapevine downy mildew is caused by the oomycete *Plasmopara viticola* (Berk. & M.A. Curt.) Berl. & De Toni, an obligate biotroph that infects grapevine species. Native to North America, this pathogen has spread to all major grapevine-growing regions worldwide, where it causes significant yield and economic losses (Toffolatti et al., 2018). In Europe, *P. viticola* was introduced in the 1870s from its center of origin (Fontaine et al., 2021, 2013; Millardet, 1881). At that time, the grapevine varieties (*Vitis vinifera*) that were cultivated there were highly susceptible to the pathogen, leading to a severe outbreak of the disease and widespread crop devastation (Kamoun et al., 2015).

The discovery of resistance sources within the *Vitis* genus led to breeding efforts that aimed at introgressing *Rpv* loci — genomic regions responsible for grapevine downy-mildew resistance— into susceptible *V. vinifera* varieties. To date, more than 30 *Rpv* loci have been identified (Possamai and Wiedemann-Merdinoglu, 2022) but only a few have been effectively incorporated into breeding programs to develop resistant grapevine varieties currently used in European vineyards. Among those, four loci —*Rpv1*, *Rpv3.1, Rpv10*, and *Rpv12*— are the most widely used and prevalent in resistant grapevine crops (Merdinoglu et al., 2018).

The genetic basis of these *Rpv* loci involves major quantitative trait loci (QTL) that exhibit monogenic inheritance pattern and mediate partial resistance phenotypes (Eibach et al., 1989), meaning that individually they are able to limit the growth and reproduction of the pathogen, but not entirely, and when combined, their inhibitory effect increases. Moreover, these loci follow a gene-for-gene interaction model, which is characterized by strain-specific responses on the same host and by a hypersensitive response when the host’s immune system recognizes the pathogen.

Despite the limited deployment of resistant varieties, *P. viticola* has already overcome *Rpv* loci-mediated resistance. The first documented breakdown resistance was reported on *Rpv3.1* by Peressotti et al. (2010)). The erosion of *Rpv10*-mediated resistance, was also reported by several studies, adaptation to *Rpv10*-carrying varieties was early spotted by Delmas et al. (2016), and later results obtained by Heyman et al. (2021) and Paineau et al. (2022) confirmed the resistance breakdown. For the *Rpv12* locus, breakdown of resistance was reported by Wingerter et al. (2021) and Paineau et al. (2022). Virulence against *Rpv* loci can be recombined into multivirulent pathotypes, and *P. viticola* strains breaking down the resistance conferred by up to two loci at a time have been described in Europe (Gouveia et al., 2024; Paineau et al., 2022; Wingerter et al., 2021). So far, *Rpv1* was the last of the main *Rpv* loci for which no breakdown had been reported, however, in 2023 an outbreak was spotted in Cilaos, Réunion Island (I. Hoarau, Personal communication, 2023), and in 2024 in France in the Rhône valley (Pelissier et al., 2025).

In recent years, gene-for-gene interactions between grapevine resistance loci and *P. viticola* effectors have been progressively deciphered. On the plant side, it has been observed that the *Rpv* loci are generally located in genomic regions that are rich in nucleotide binding domain and leucine-rich repeat genes (NBS-LRR) which are involved in the recognition of specialized pathogen effectors (Di Gaspero et al., 2007; Merdinoglu et al., 2018; Moroldo et al., 2008). On the pathogen side, the RxLR family is the largest and most studied family of cytoplasmic effectors in oomycetes. These effectors are characterized by the presence of a signal peptide (SP), a RxLR-EER motif at their N-terminal sequence with one or more WY-domains, and a common fold found only in this family of proteins (Anderson et al., 2015).

More specifically, loci associated with the virulence of *P. viticola* on *Rpv3.1*, *Rpv10* and *Rpv12* have been identified through Genome Wide Association Studies (GWAS) (Paineau et al., 2024) and Quantitative Trait Loci (QTL) mapping (Dvorak et al., 2025a). Virulence towards *Rpv3.1* and *Rpv12*-carrying varieties was found to be recessive and associated with deletions or disruption of genes encoding RxLR effector-like proteins (Dvorak et al., 2025a; Paineau et al., 2024). Outbreaks associated with *Rpv3.1* and *Rpv12* breakdown appear to result from recurrent, independent mutation events across different geographical regions (Delmotte et al., 2014; Dvorak et al., 2025a; Paineau et al., 2024). The locus involved in the *Rpv10* breakdown involved a different underlying mechanism, the virulence was dominant and linked to the presence of additional putative secreted proteins, suggesting a possible virulence-suppressor mechanism (Dvorak et al., 2025a). The haplotype carrying the allele conferring virulence towards *Rpv10* varieties seemed to have originated from *P. viticola* north-American populations and spread into some European areas via secondary introduction of the pathogen (Dvorak et al., 2025a).

In this study, we investigated the grapevine downy mildew outbreak detected in 2023 in Cilaos, where a marked increase in disease incidence was observed in vineyards planted with the ResDuR variety Floreal, which carries *Rpv1* and *Rpv3.1* (I. Hoarau, Personal communication, 2023). We isolated and characterized a strain, referred hereafter as the “Cilaos strain”, assessing its virulence profile and genetic background together with a panel of European strains representing *P. viticola* genetic and virulence diversity. Whole-genome sequencing of these strains was used to assess the genetic origin of the Cilaos strain and characterize the alleles at the known avirulence loci (Dvorak et al., 2025a; Paineau et al., 2024). Our results provide the first evidence for the breakdown of *Rpv1* locus. Moreover, this represents the first documented case of a *P. viticola* strain able to simultaneously overcome three out of the four mainly deployed *Rpv* loci (*Rpv1*, *Rpv3.1* and *Rpv10*). Remarkably, the Cilaos strain exhibited an unprecedent complete loss of host recognition on *Rpv10*-carrying varieties and harbored a novel mutation at the avirulence locus, distinct from the previous breakdown of *Rpv10* in Europe.

## Materials and Methods

### Virulence profile characterization

#### Plasmopara viticola strains

A strain originating from Cilaos on Réunion Island (Pv8586_1), was obtained through isolation as described by Paineau et al. (2022), from a vineyard where increased virulence was observed on the Floreal variety, which carries the *Rpv1* and *Rpv3.1* loci (*Rpv1*-*Rpv3.1*). In addition, eight other strains previously obtained in separate studies (Paineau et al., 2022; Wingerter et al., 2021) and originating from various European wine-producing countries were included in this study due to their diverse virulence profiles against *Rpv* loci (see Table 1). Of these eight strains, five (Pv1419_1, Pv2543_1, Pv2664_1, Pv412_11, and Pv8522_1) were previously shown to overcome resistance conferred by some of the most commonly used *Rpv* loci, *Rpv3.1*, *Rpv10*, and *Rpv12*. In contrast, the remaining three strains (Pv2578_1, Pv2909_11, and Pv3174_11) were isolated from grapevine varieties lacking any known *Rpv* loci and appear to be non-virulent on *Rpv*-carrying varieties (Table 1).

**Table 1.** Panel of *Plasmopara viticola* strains with variated origins and virulence profiles, and their associated genomic data availability.

| Strain ID | Reported virulence | Year of collection | Country | Locality | Reference | Genomic data Accession number |
| --- | --- | --- | --- | --- | --- | --- |
| Pv8586_1 | - | 2023 | France | Cilaos-Réunion Island | - | PRJNA 1255592 |
| Pv2664_1 | *Rpv3.1* | 2013 | France | Villenave d'Ornon | Paineau et al. (2022) | PRJNA 1255592 |
| Pv412_11 | *Rpv3.1* | 2010 | Switzerland | Cugnasco | Paineau et al. (2022) | PRJNA 1095879 |
| Pv8522_1<sup>a</sup> | *Rpv3.1/Rpv12* | NA | Switzerland | Soyhières | (Wingerter et al., 2021) | PRJNA 875296 |
| Pv1419_1 | *Rpv3.1/Rpv10* | 2012 | Germany | Ihringen | (Paineau et al., 2022) | PRJNA 1095879 |
| Pv2543_1 | *Rpv12* | 2013 | Hungary | Pécs | Paineau et al. (2022) | PRJNA 1095879 |
| Pv2578_1 | Avirulent | 2013 | France | Le Landreau | (Paineau et al., 2022) | PRJNA 1255592 |
| Pv3174_11 | Avirulent | 2016 | Italy | Piateda | (Paineau et al., 2022) | PRJNA 644583 |
| Pv2909_11 | Avirulent | 2014 | Germany | Geisenheim | Paineau et al. (2022) | PRJNA 1255592 |

<sup>a</sup> Name of the strain in Wingerter et al. (2021) publication was “VB-isolate”
NA: no information

### Plant material

To assess the virulence of the Cilaos strain together with the other chosen strains, a cross-inoculation experiment was performed in which four grapevine varieties carrying different resistance loci were used. i.e. 3179-90-7 (*Rpv1*), Regent (*Rpv3.1*), Muscaris (*Rpv10*) and Fleurtai (*Rpv12*). Additionally, the susceptible variety *Vitis vinifera* cv. Chardonnay that carries no resistance loci was included as a control, for a total of five varieties.

A total of 20 plants per each variety were collected from nurseries and grafted onto the Selection Oppeinheim 4 (SO4) rootstock. All the plants were grown simultaneously in a glasshouse under natural photoperiod conditions, without chemical treatment. After two months of growth, the 5th leaf below the apex was collected and used for the inoculation.

### Cross-Inoculation assay

Inoculation was carried out according to the methodology of Paineau et al. (2022), with modifications. To obtain the inoculum, strains were propagated on leaves of *Vitis vinifera* cv. Cabernet Sauvignon. The day before inoculation, sporangia were gently rinsed off to ensure that freshly produced, uniformly aged sporangia could be collected the following day. The day of the inoculation, a sporangia suspension was prepared and adjusted to a concentration of 1×10⁵sporangia/mL using a portable particle counter (Scepter 2.0 Automated Cell Counter; Millipore).

The nine strains were inoculated on the five grapevine varieties, resulting in a dataset of 45 interactions. For each interaction, six repetitions were performed as described below. Inoculation was carried out on 15mm diameter leaf discs excised from the fifth fully expanded leaf below the apex of each plant. To prepare the discs, leaves were first washed with reverse-osmosis water, dried on absorbent paper, and then the discs were excised using a cork borer. For each interaction, six discs (representing repetitions) from different leaves were placed abaxial side up on wet filter paper within a Petri dish. All the discs that were going to be inoculated with one strain were randomly arranged within the dish. This represented a total of 30 discs, per Petri Dish, sprayed with 4mL of the sporangia suspension. Mock inoculations were also performed using sterile water instead of the sporangia suspension.

In summary, six repetitions were conducted for each combination of variety and strain, resulting in a total of 240 inoculated discs for the experiment, plus the additional 30 mock-inoculated discs.

One day after inoculation the leaf discs were wipe-dried with filter paper. The Petri dishes were sealed with plastic film to maintain high relative humidity and kept under controlled conditions during six days at 18°C under a 12/12h photoperiod.

After performing this cross-inoculation assay, a second assay was performed to confirm results. In this experiment only the varieties Chardonnay (susceptible), 3179-90-7 (*Rpv1*) and Regent (*Rpv3.1*) were inoculated with all the strains except Pv412_11. Additionally, the assay was replicated using the fifth, eight and tenth fully expanded leaves below the apex of each plant.

### Symptoms and virulence assessment

Sporulation and necrosis were measured at the 6th day post-inoculation on all the leaf discs used. Sporulation area was quantified by taking high resolution photos and then performing an image analysis to estimate the proportion of leaf area covered by sporangia. Each photo included four leaf discs. To take the pictures we used a Canon EOS 650D camera equipped with a macro lens (Canon EF 100 mm f/2.8 USM). Photographs were taken in manual mode (f/5.6; ISO-100). Each 17.9-megapixel image was analyzed with the Python-based tool CATIA (available at: https://gitlab.com/grapevinedownymildew/notebook_image_analysis) trained to isolate the discs and then detect the sporulation area.

Necrosis was assessed as an indicator of the hypersensitive response (HR) using an adaptation of the visual scale proposed by Paineau et al. (2022, Fig. S1), with the addition of a value of 0 to indicate the absence of necrosis. We developed a visual scale ranging from 0 to 5, where 0 and 1 correspond to the absence of an HR, and values from 2 to 5 indicate the presence of HR in the host variety in different categories.

The different types of necrosis were classified based on their shape and distribution across the leaf, as illustrated in Figure 1, to reflect the type of necrosis involved in the plant’s immune response.

**Figure 1.**
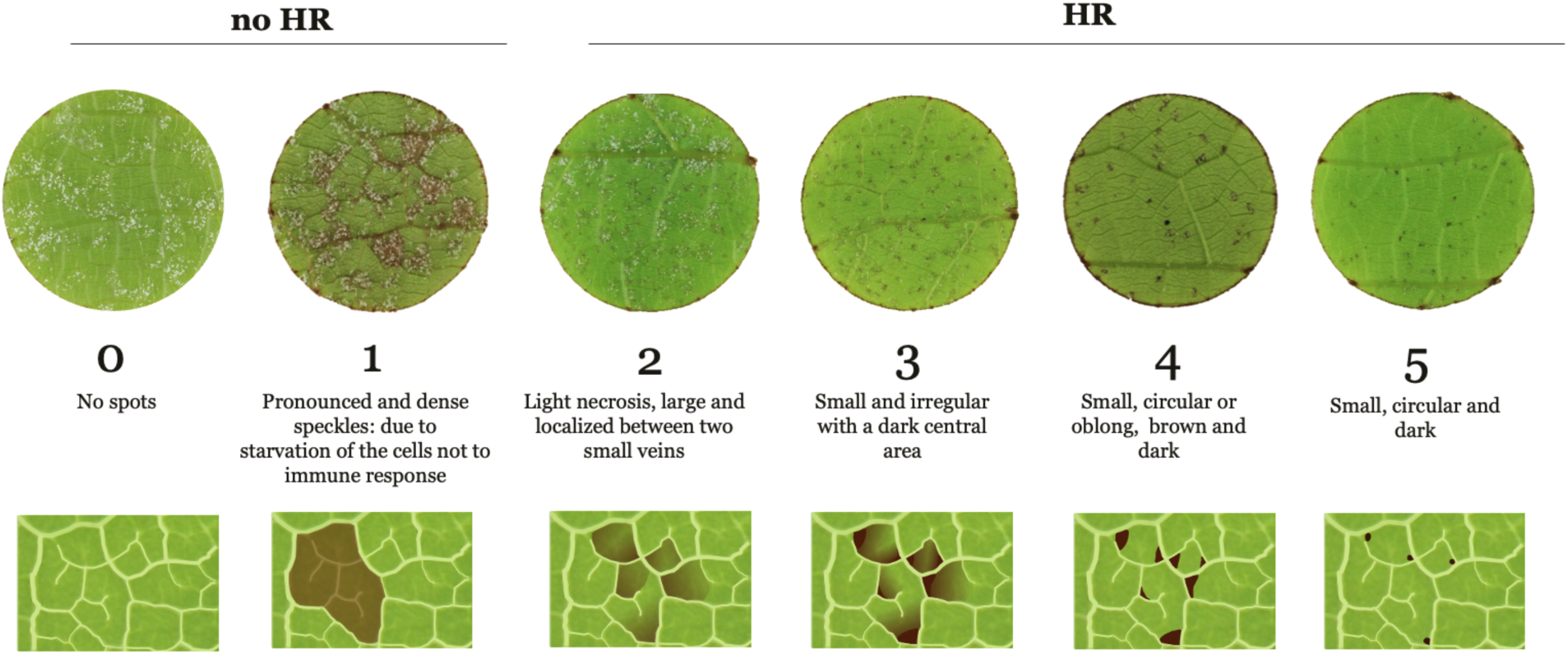
Necrosis visual scale used to asses Hypersensitive Response (HR) to *Plasmopara viticola* infection on grapevine leaf discs that were spray-inoculated. On the top of the figure is displayed the classification of the different values of the scale according to the absence (**no HR**) or presence of HR (**HR**). Pictures of infected leaf discs illustrating the different categories of the scale are displayed with a brief description of the different values of HR according to the observed necrosis pattern. In the bottom of the figure there is the corresponding graphical representation of the different categories of HR. **0**: no necrosis nor visible tissue damage, **1**: Dense brownish lesions that extend without clear delimitation, typical of tissue damage due to cell starvation but not caused by immune response of the plant **2**: Light diffused necrosis spots localized inside the limits of the polygons formed by the small veins of the leaf, **3**: Smaller necrosis spots, still inside the limits created by the small veins of the leaf, with a central point that diffuses inside those limits, **4**: Small circular or oblong spots that do not diffuse and localize in the corners formed by the intersection of the small veins of the leaf, **5**: Very small and well defined dots that tend to be circular and localize in the intersection of the small veins of the leaf.

Additionally, we used a parameter that we called Efficiency of the Resistance Factor (ERF) to assess the capacity of a given strain to sporulate on a variety carrying a given *Rpv* locus in comparison with the capacity to sporulate on a variety with no resistance locus, i.e. susceptible, following:

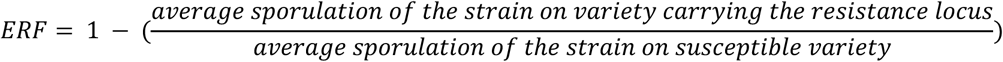

The value is expressed as the complementary ratio between the sporulation of the strain on the variety carrying the resistance locus and the sporulation of the strain on a susceptible variety, as defined in Formula 1. The lower the value of the ERF, the higher the sporulation on the locus-carrying variety having as reference the sporulation on the susceptible variety. Negative values can be obtained when sporulation on the *Rpv*-carrying variety is higher than on the susceptible variety.

We considered that a strain was able to breakdown the resistance conferred by a given locus similarly to the method described by Paineau et al., (2022), based on the principle that a virulent strain should be able to evade the host’s immune system, i.e. not producing HR, and being able to propagate, i.e. have efficient spore production on both susceptible and *Rpv*-carrying variety.

### Characterization of the *Plasmopara viticola* loci *AvrRpv3.1,* S-*AvrRpv10* and *AvrRpv12* in the Cilaos strain

To further characterize the Cilaos strain, we analyzed the previously identified *P. viticola* loci that interact with the grapevine resistance loci *Rpv3.1*, *Rpv10*, and *Rpv12*. (Dvorak et al., 2025a; Paineau et al., 2024). This analysis included DNA extraction from the Cilaos strain, sequencing, and mapping of the reads—together with publicly available reads from the other phenotyped strains—against reference genomes. The accession numbers for the sequencing reads available in public databases are listed in Table 1.

To obtain the DNA from the Cilaos strain, sporangia, and sporangiophores were produced and collected following the protocol described by Dussert et al. (2019). DNA extraction was performed using a CTAB-based protocol adapted from Möller et al. (1992) and further modified by Dvorak et al. (2025b). The purity and concentration of the extracted DNA were assessed using a DeNovix DS-11 spectrophotometer and a Qubit 3 fluorometer (Invitrogen), respectively. Libraries were prepared using an Illumina DNA TruSeqkit and sequencing was performed at the GeT-PlaGe facility (Toulouse, France https://get.genotoul.fr/) with a NovaSeq6000 to produce 2×150 bp paired-end read. Reads from all phenotyped strains were mapped to different reference genomes using BWA MEM2 v2.2.1 (Li, 2013). For the *P. viticola* locus associated with the *Rpv3.1* breakdown, we used the reference genome Pv221_FU (Paineau et al., 2024) (genome available: https://doi.org/10.57745/MXJWZS). The interval between 640 and 740 kbp of the Primary_000014F contig was then analyzed using the corresponding genome annotation as guide. For the locus associated with *Rpv10*, we used the diploid genome assembly Pv1419_1 and its gene annotation (Dvorak et al., 2025b), examining both haplotypes, as this region appears to have undergone substantial rearrangements and the gene content differ between them. In Haplotype 1, we examined the region of chromosome 16 (Chr16) spanning 3,900–4,400 kbp. In Haplotype 2, which is known to harbor the virulent allele, the analyzed region was on chromosome 16, but between 4,050–4,750 kbp. Finally, for the locus associated with *Rpv12*, we used the same reference genome as for *Rpv3.1*, but analyzing the region on the Primary_000017 contig from 250 to 400 kbp.

Afterward, we computed both the average genomic depth and the depth across sliding windows within the target loci to obtain normalized read depth. This allowed us to estimate gene copy number as a proxy for ploidy and to identify allele variation at each locus. The sliding window sizes used for the genomic regions associated with *Rpv3.1*, *Rpv10*, and *Rpv12* were 1kbp, 4kbp, and 1kbp respectively. To identify nonsense mutations and any open reading frame disruptions, mapping depths and alignments within the targeted loci were visualized using IGV V2.19 (Robinson et al., 2011).

Accessions numbers of all sequencing data used in this analysis are listed in Table 1.

### Mating type

To further characterize the Cilaos strain and the other strains in the panel, mating type was determined through crosses with reference strains of known mating types P1 and P2, following the method described by Dussert et al. (2020). In summary, each strain was first grown on grapevine leaves, a sporangial suspension was then prepared for each strain and mixed with a sporangial suspension of either the P1 or P2 reference strain. The mixtures were inoculated onto grapevine leaf discs placed on water agar and incubated at 22 °C for 5 days (12 h/12 h photoperiod), after which the temperature was lowered to 12 °C and maintained for three weeks. After the incubation period, the presence of oospores was assessed using a binocular magnifier. A strain that produced oospores only with one of the reference strains was classified as the opposite mating type of that reference.

### Population structure analysis

To understand the genetic relationships of the Cilaos strain within the group of strains, a population structure analysis was carried out. To genotype the strains, the publicly available reads from whole genome sequencing were mapped against the reference genome Pv221_FU using BWA MEM2 v2.2.1 (Li, 2013) and bam files were handled using SAMTOOLS v1.10 (Li et al., 2009). The PCR duplicates were removed with the tool MARKDUPLICATES in GATK v 4.2.6.1 (McKenna et al., 2010). Variant calling was performed in the same software with the tool HAPLOTYPECALLER setting the option output-mode to EMIT_ALL_ACTIVE_SITES. This procedure was performed parallelizing mapping and variant calling using the RATTLESNP pipeline (https://rattlesnp.readthedocs.io/).

After the variant calling step, high-quality sites were filtered using BCFTOOLS v1.21 (Li, 2011) and VCFTOOLS V0.1.16 (Danecek et al., 2011) to obtain a dataset with the following characteristics: indels were removed, only biallelic SNPs (Single Nucleotide Polymorphisms) were kept, SNPs in repetitive regions were removed, and SNPs were kept when MQ ≥ 59, QUAL ≥ 25, mean depth ≥ 5 and ≤ 70, then genotypes at each site with depth values outside the aforementioned range were set to missing, and finally no missing data per site was allowed.

The resulting high-quality SNPs were used to analyze the placement of the Cilaos strain (Pv8586_1) into the group of *P. viticola* strains. For this, we performed a Principal Components Analysis (PCA) using the ADEGENET package (Jombart and Ahmed, 2011) in R 4.4.1 and used the clustering algorithm SNMF (sparse non-negative matrix factorization) implemented in the package LEA (Frichot et al., 2014) in the same software.

An additional analysis was done with a broader panel of strains previously presented by (Dvorak et al., 2025a) to confirm results (list of strains in Supplementary table 1).

## Results

### The Cilaos strain is able to break down the resistance conferred by *Rpv1* and other *Rpv* loci

In the first inoculation experiment five varieties were inoculated with the Cilaos strain and the other eight strains. All strains successfully infected the susceptible variety Chardonnay, evading host’s immune response (no HR was observed) and displaying high values of sporulation, as shown in Figure 2A. Control mock inoculated disks remained healthy until the end of the experiment.

**Figure 2.**
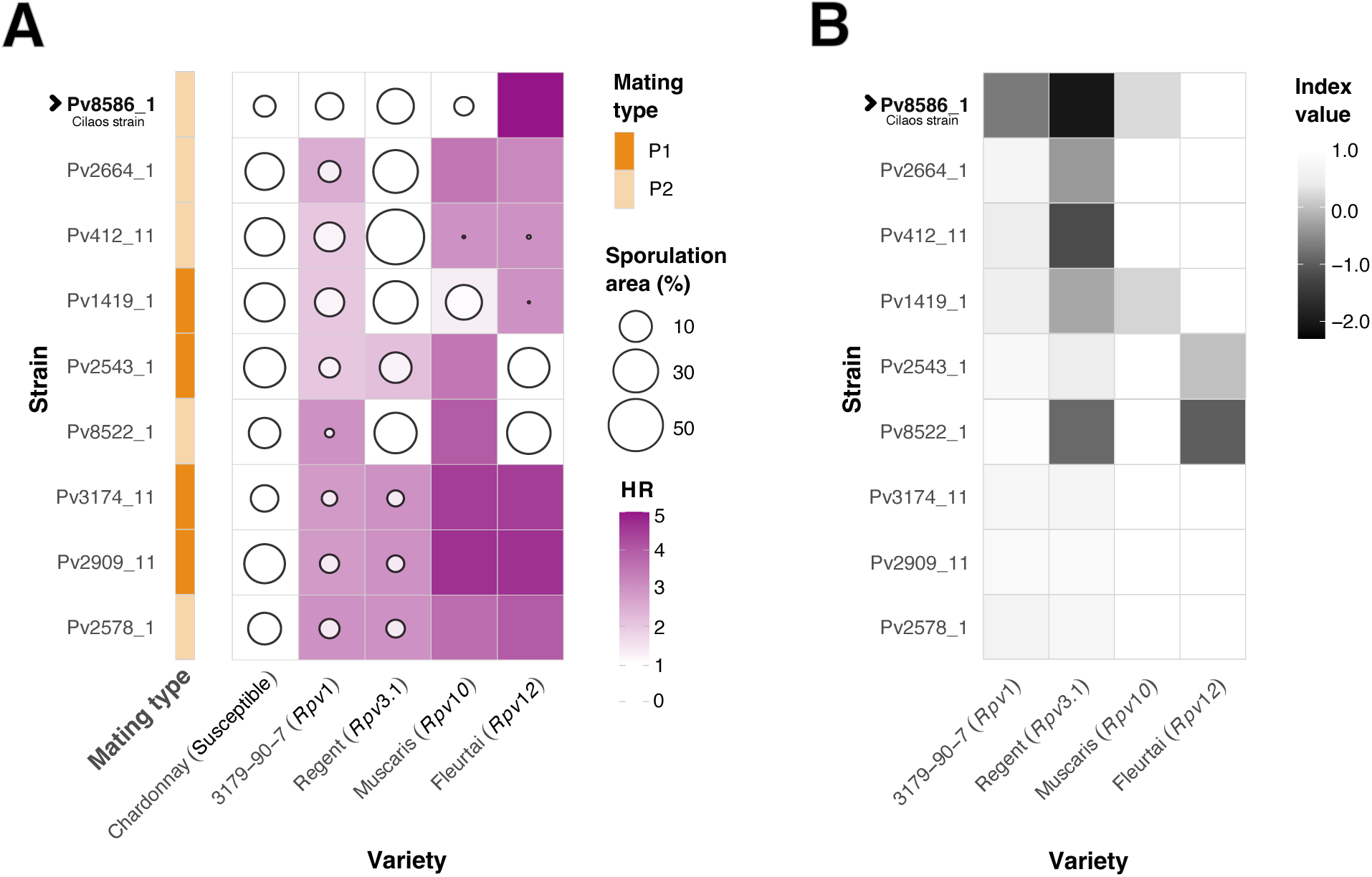
Mating type and virulence of the *Plasmopara viticola* strain Pv8586_1 from Cilaos. Eight extra strains with different virulence profiles and geographical origins were included as reference. Results of a cross-inoculation experiment are displayed, in which all strains were inoculated on different varieties carrying the loci *Rpv1*, *Rpv3.1*, *Rpv10* and *Rpv12* and on a susceptible variety with no *Rpv* loci (Chardonnay). **A.** The strain names are displayed on the Y axis alongside a colored bar indicating their mating type. Adjacent to the bar, a heatmap presents the percentage of sporulating area and the Hypersensitive Response (HR) score for each interaction between Variety and Strain, varieties are shown in the X axis. For the HR, the heatmap shows the mean value of the five repetitions, represented on a color scale. For the sporulating area, values are represented by circles, with the size of each circle corresponding to the mean of the percentage value. **B.** Heatmap that shows the values of Efficiency of the Resistance Factor (ERF index) for each interaction tested, except for the susceptible variety.

As expected, the strains Pv3174_11, Pv2909_11, and Pv2578_1 exhibited an avirulent phenotype, being unable to evade the immune response of grapevine varieties that carry *Rpv* loci. Sporulation was either absent or substantially lower on varieties with the *Rpv1* and *Rpv3.1* loci compared to the susceptible variety. As for the rest of the panel of strains, Pv2664_1, Pv412_11, Pv1419_1, Pv2543_1 and Pv8522_1, they were shown to be virulent and met our definition of resistance breakdown, displaying no HR and efficient sporulation on the *Rpv* loci on which they were reported to be virulent (Figure 2, Supplementary table 2) (see reported virulence in Table 1). The only exception was strain Pv1419_1 which breaks down the resistance conferred by *Rpv3.1* but only partially breaks down *Rpv10*. It was able to sporulate efficiently on the *Rpv10*-carrying variety displaying a low value of ERF (ERF value = 0.1), but did not evade HR in all inoculated discs of this variety. Overall, these results aligned with the known virulence profiles of the tested strains and confirmed the reliability of the assay.

For the Cilaos strain (Pv8586_1), this assay showed that this strain is able to breakdown *Rpv1*-mediated resistance consistent with the field observations on Réunion Island. It further revealed that the strain was also able to overcome *Rpv3.1* and, most notably, *Rpv10*. When inoculated on varieties carrying *Rpv1*, *Rpv3.1*, or *Rpv10*, the Cilaos strain did not trigger a HR in any of the leaf discs and sporulated abundantly, resulting in low ERF values (-0.6,-2 and 0.2 respectively) (Figure 2, Supplementary table 2), and therefore, meeting our definition of a breakdown for these three loci.

Interestingly, when inoculating strains that broke down a specific *Rpv* locus on the variety carrying that locus, the ERF value was always below 0.3 (Supplementary table 2).

Results from the second inoculation experiment confirmed the virulence profiles of the Cilaos strain on *Rpv1* and *Rpv3.1* (Supplementary Figure 1, Supplementary table 3).

To further characterize the Cilaos strain, along with the other strains used in the inoculation experiments, we assessed the mating type through crossing tests. The Cilaos strain was classified as P2; four additional strains were also classified as P2, while the remaining strains belonged to the P1 type (Figure 2A).

### Characterization of the *Plasmopara viticola* loci *AvrRpv3.1,* S-*AvrRpv10* and *AvrRpv12* in the Cilaos strain

To further characterize the Cilaos strain, we examined genomic regions and their gene content at *P. viticola* loci previously associated with the breakdown of *Rpv3.1*, *Rpv10*, and *Rpv12* resistance, analyzing normalized mapping depths (Supplementary Figures 2–5), and compared these results with those from the other eight strains used as references.

For the locus associated with *Rpv3.1* interaction, the Cilaos strain Pv8586_1 showed a complete absence of the genes encoding effectors as expected for a virulent genotype (Figure 3, Supplementary Figure 2) and was shown to carry a virulence allele previously described as *avr1* by Paineau et al. (2024). All strains that were virulent on *Rpv3.1*-carrying varieties, including the Cilaos strain, displayed either a complete absence, partial deletions, or nonsense mutations in these effectors (Figure 3).

**Figure 3.**
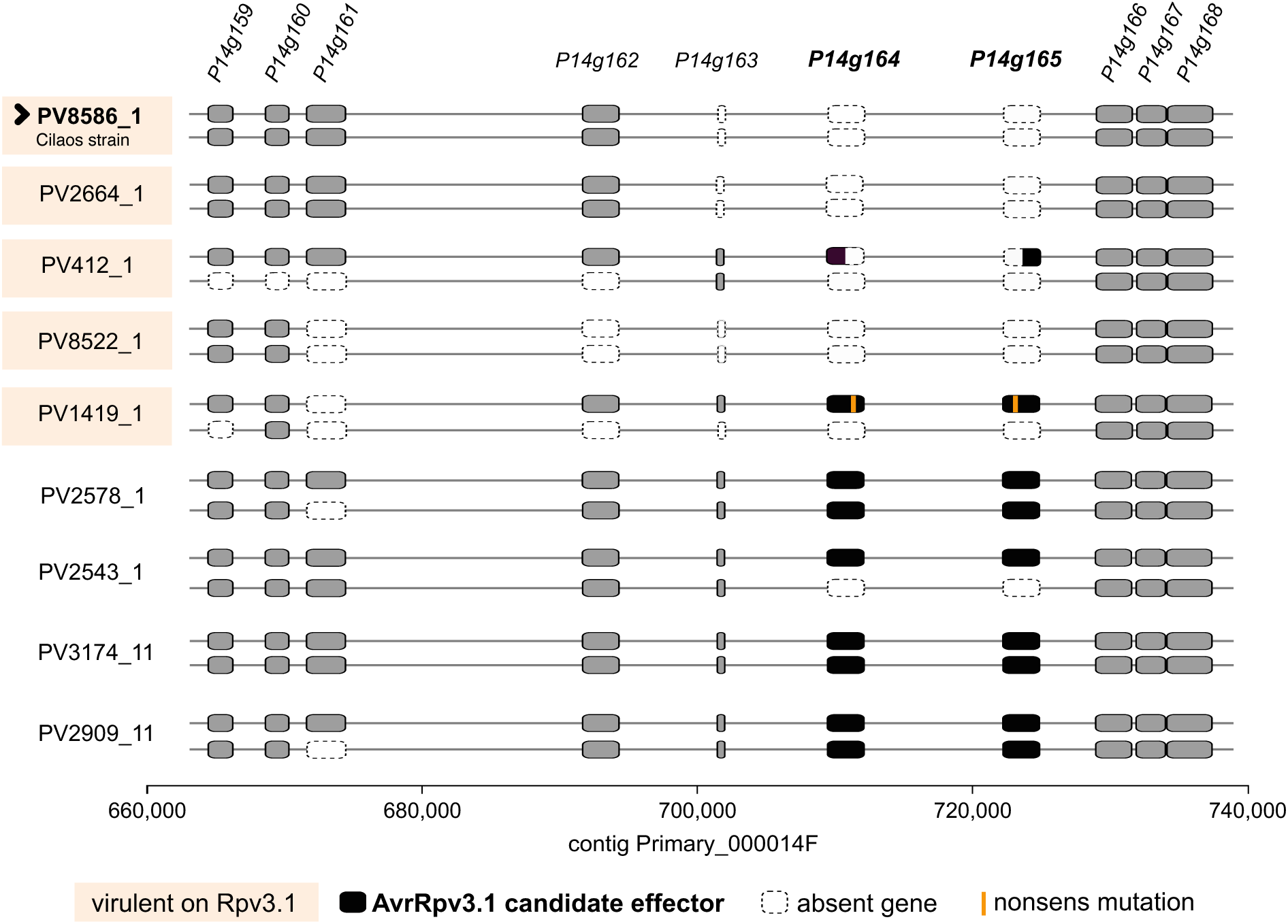
*Plasmopara viticola* locus associated with *Rpv3.1* breakdown in a Cilaos strain and eight additional strains of variated virulence profiles. Strains that are virulent on *Rpv3.1*-carrying grapevine varieties are highlighted in light peach. For each strain the locus is represented by two lines that correspond to the haplotypes (*P. viticola* is diploid), with genes shown as boxes along the sequence, gene names are shown above each graphic. Black boxes indicate candidate-effector genes. Gray boxes denote other genes, while dashed white boxes indicate gene absence. Nonsense mutations in genes are shown in bright orange. The Cilaos strain is marked with an arrow and bold letters.

To study the characteristics of the genomic region associated with *Rpv10* resistance breakdown, and to take into account the complexity of the genomic architecture of the region, we used both haplotypes of the partially virulent strain Pv1419_1, in which the *Rpv10* virulence mechanism was originally identified, as reference. These were compared to the corresponding region in the fully virulent Cilaos strain and in the remaining strains of the panel, all of which are avirulent on *Rpv10*.

In the strain used as reference we cand find the haplotypes previously described by Dvorak et al. (2025a): One that is present in avirulent European populations (Haplotype 1), and the other (Haplotype 2), which is thought to be inherited from American populations and confers virulence on *Rpv10* by a dominant suppressor mechanism. The presence or absence of these haplotypes, as well as their genes encoding secreted proteins, including the candidate suppressors (S-*AvrRpv10*), was assessed (Supplementary Figure 3 and 4). We found that all avirulent strains lacked the American allele and the putative suppressors, as expected (Figure 4). However, the Cilaos strain also lacked this allele despite displaying a virulent phenotype on *Rpv10*-carrying varieties. Instead, this strain harbors a homozygous large deletion (∼120–215 kb) spanning most of the examined region (Figure 4, Supplementary Figure 3 and 4).

**Figure 4.**
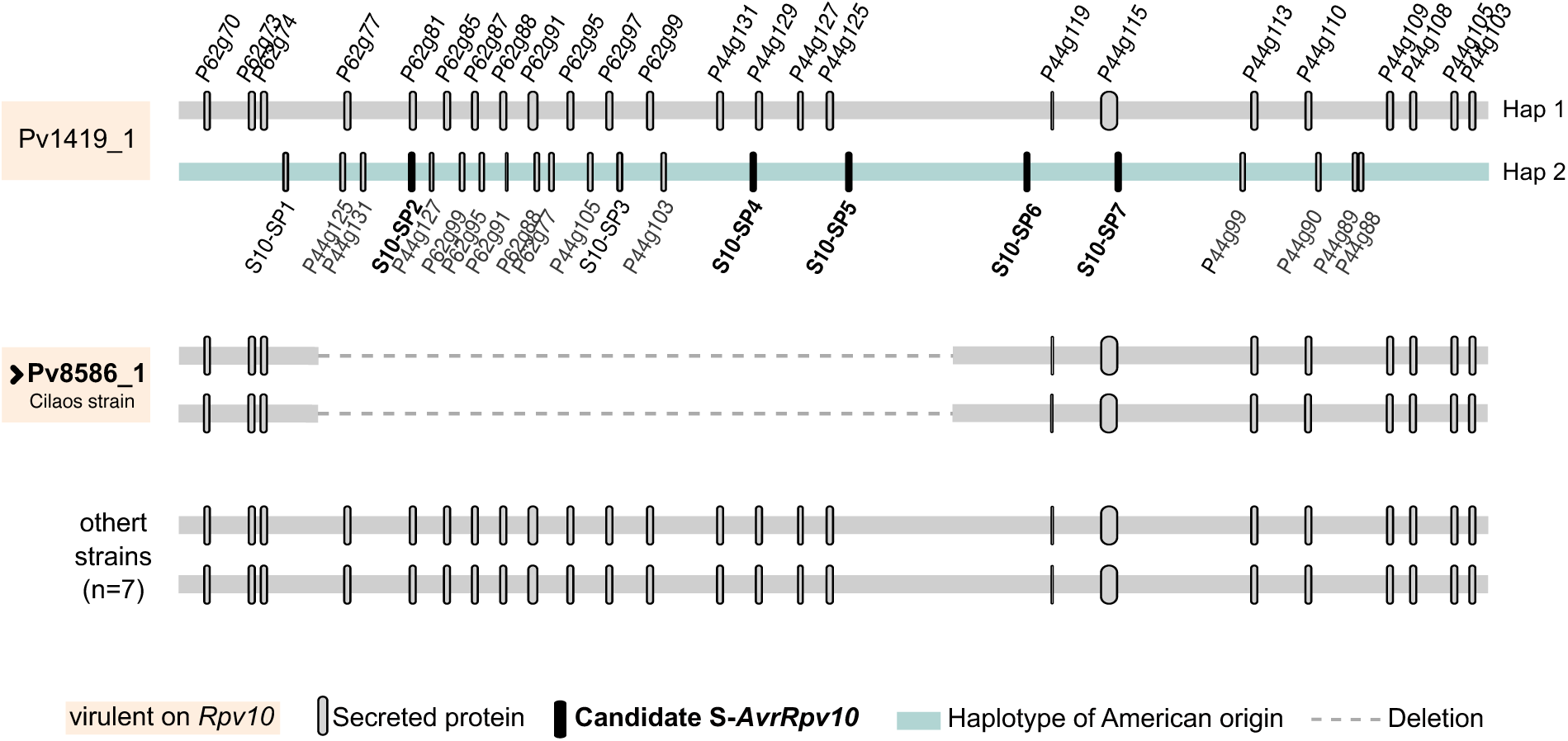
Comparison of the *Plasmopara viticola* genomic region associated with virulence on *Rpv10* among a reference virulent strain (Pv1419_1), a virulent strain from Cilaos - Réunion Island (Pv8586_1), and seven avirulent strains (Pv3174_11, Pv2909_11, Pv2578_1, Pv2664_1, Pv412_11, Pv2543_1 and Pv8522_1). The figure shows a non-scaled diploidized representation of the associated genomic region. For each strain or strain group, each band corresponds to a haplotype sequence (*P. viticola* is diploid); dashed lines indicate deletions or absence of sequence. Genes encoding secreted proteins are shown as ellipsoidal gray boxes, with black boxes representing candidate virulence suppressors (S-*AvrRpv10*). At the top of the graphic the haplotypes of the virulent European strain Pv1419_1 described by Dvorak et al. (2025a) https://doi.org/10.1101/2025.05.18.654733) are displayed. Hap 1 represents the European haplotype found in avirulent strains across Europe, whereas Hap 2 corresponds to the rearranged haplotype of American origin that carries putative virulent suppressors and that is thought to confer virulence on *Rpv10* by dominant inheritance. The American-derived allele is shown in pale green. Below, the Cilaos strain is shown, characterized by a homozygous deletion in the region and absence of the candidate suppressors and the American allele. The rest of the strains (other strains) show no modification in this region and do not carry the American allele and the candidate suppressors. Strains that are virulent on *Rpv10*-carrying grapevine varieties are highlighted in light peach. The Cilaos strain is marked with an arrow and bold letters.

Finally, for the *AvrRpv12* locus, the strains that were virulent on *Rpv12* lacked the putative effectors linked to avirulence (Pv8522_1 and Pv2543_1). In contrast, the Cilaos strain, along with the other strains that were avirulent on the *Rpv12*-carrying variety, had the putative effectors (Figure 5, Supplementary figure 5).

**Figure 5.**
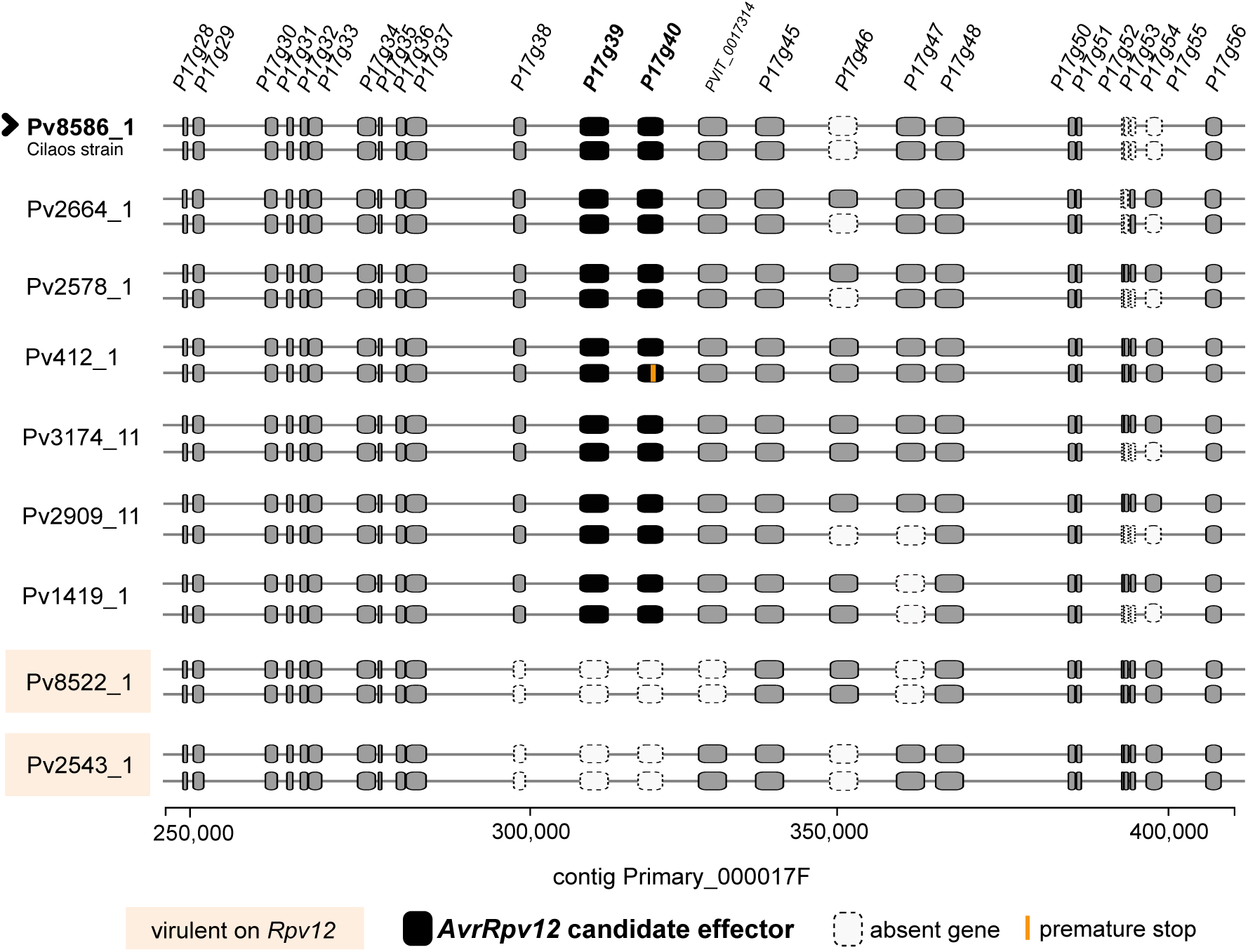
*Plasmopara viticola* locus associated with *Rpv12* resistance breakdown in a Cilaos strain and eight additional strains of variated virulence profiles. Strains that are virulent on *Rpv12*-carrying grapevine varieties are highlighted in light peach. For each strain the locus is represented by two lines that correspond to the haplotypes, with genes shown as boxes along the sequence, gene names are shown above each graphic. Black boxes indicate candidate-effector genes. Gray boxes denote other genes, while dashed white boxes indicate gene absence. Premature stop codon in genes is shown in bright orange. The Cilaos strain is marked with an arrow and bold letters.

### The Cilaos strain is related to French and other European strains

After variant calling and quality filtering, a total of 257,681 high-quality biallelic SNPs were retained. This dataset was then used for a population structure analysis.

A principal component analysis (PCA) was performed, and the plot of PC1 versus PC2 (Figure 6A) showed that the Cilaos strain (Pv8586_1) clustered closely with most continental European strains, except with Pv1419_1, one of the included strains originating from Germany. To better assess the relationship between the Cilaos strain and the other strains, PC2 and PC3 were also examined (Figure 6B). This analysis revealed that the strain most closely related to the Cilaos strain was Pv2664_1 from continental France (Figure 6B).

**Figure 6.**
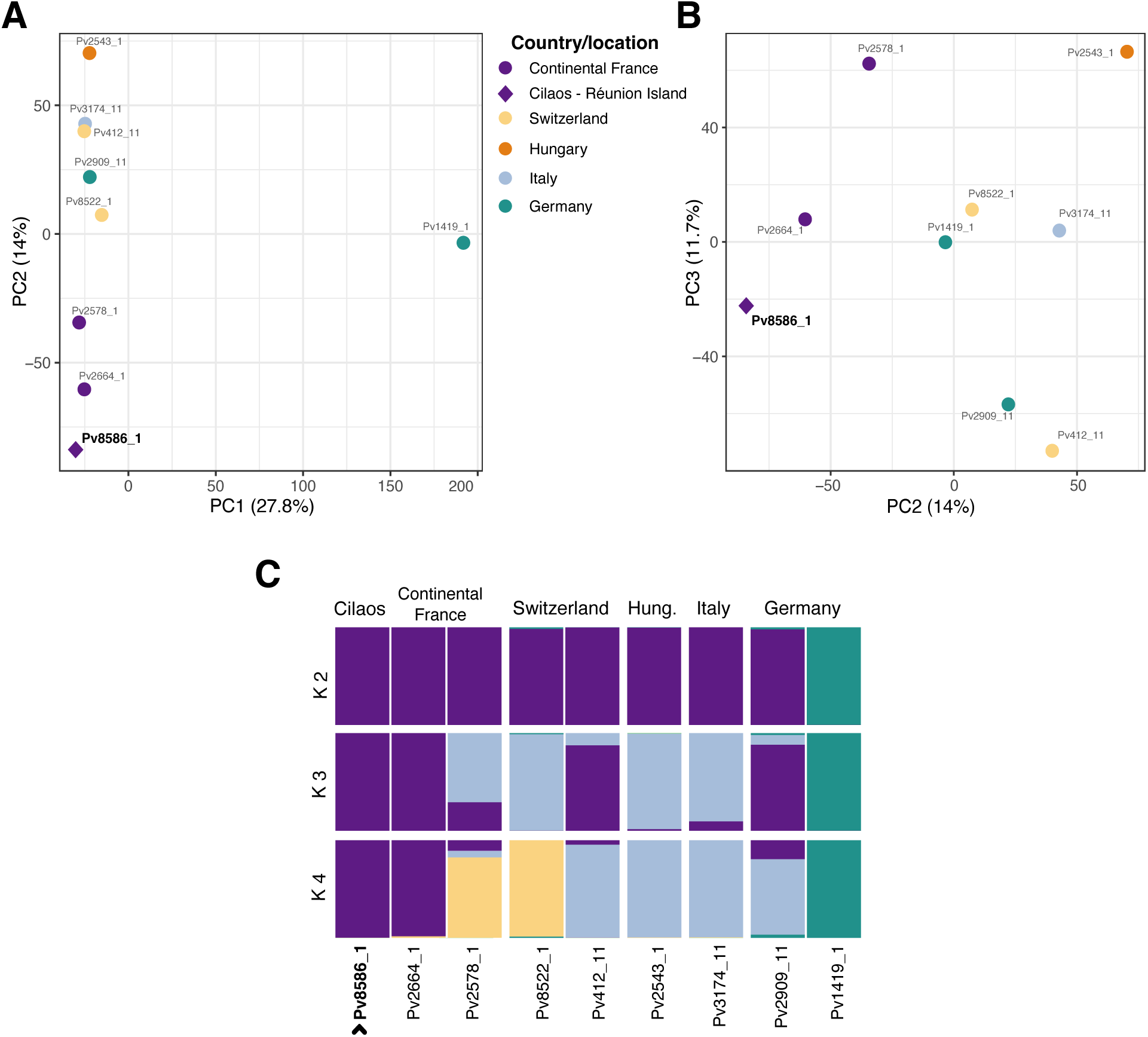
The *Plasmopara viticola* strain from Cilaos (Réunion Island, France) that breaks down *Rpv1* mediated resistance is genetically similar to European strains and closely related to a continental French strain. Analyses were performed including nine *P. viticola* strains and using 257,681 high quality SNPs (Single Nucleotide Polymorphism). **A**. Principal Components Analysis (PCA) of the SNPs from *P. viticola* strains. PC1 and PC2 are displayed along with the variance explained by each. Country of origin per each strain is shown. Cilaos strain is highlighted in bold letters. **B**. PCA of the genetic variants (SNPs) from *P. viticola* strains. PC2 and PC3 are displayed along with the variance explained by each. Country of origin per each strain is shown following the same color code than shown in the A-part of this figure. Cilaos strain is highlighted in bold letters. **C**. Genetic clustering results inferred using the SNMF (sparse non-negative matrix factorization) algorithm. Each vertical bar represents an individual strain, labeled along the X-axis. The colors within each bar indicate the estimated ancestry proportions in given K number of populations, with K values shown on the left margin of the plot. Isolates are ordered by country. Cilaos strain is indicated by an arrow and bold letters.

The sparse non-negative matrix factorization (SNMF) algorithm was also applied. Results indicated that the Cilaos strain (Pv8586_1) was genetically closer to strains from continental France (Pv2664_1), Switzerland (Pv412_11), and Germany (Pv2909_11) (Figure 6C, K=3). Among these, the French strain Pv2664_1 was identified as the most genetically similar to the Cilaos strain (Figure 6C, K=4).

Overall, the strains show strong population differentiation, forming two distinct groups: one consisting solely of the German strain Pv1419_1, and another comprising all remaining strains. The Cilaos strain appears to share a common genetic background with the latter group, and inside this group it seems to have more genetic similarities with the French strains. These results were further supported by analysis of a broader panel of strains including non-European ones (Supplementary Figure 6).

## Discussion

Our results demonstrate that the *Plasmopara viticola* strain Pv8586_1, isolated from Cilaos on Réunion Island, is overcoming the resistance conferred by the *Rpv1* locus. The virulent phenotype of the Cilaos strain on *Rpv1*-carrying grapevine varieties is clear and reproducible: the strain produces abundant sporulation and completely escapes host recognition, as evidenced by the absence of HR in all the replicates. Population genomics analysis revealed that the Cilaos strain shares a genetic background closely related to *P. viticola* strains sampled in continental France, suggesting a likely European origin.

In addition, phenotypic assays also showed that the Cilaos strain exhibits a multi-virulent profile, being able to overcome resistance mediated by both the *Rpv3.1* and *Rpv10* loci. The breakdown of *Rpv3.1* in this strain follows the previously described mechanism (Dvorak et al., 2025a; Paineau et al., 2024), with a recessive mode of inheritance and involving the same virulence determinants. For Rpv10, however, our results indicate a novel breakdown mechanism. In this case, the loss of resistance is associated with a complete absence of pathogen recognition by the plant, distinct from the incomplete HR previously reported in Europe. Furthermore, the strain exhibited a large homozygous deletion in the corresponding *avr* locus, representing a previously unreported mutational event that could be associated with *Rpv10* breakdown. These findings revealed that *P. viticola* can evade recognition by more than two resistance genes, exceeding the previously reported maximum of two (Paineau et al., 2022), and is capable of accumulating multiple virulence factors (three in this case) within a single genotype, highlighting its unprecedented evolutionary potential to overcome a broad spectrum of major grapevine resistance loci.

The emergence of this multivirulent strain may have been facilitated by local environmental conditions in Cilaos, which are highly favorable to downy mildew development, in combination with the deployment of grapevine varieties carrying distinct *Rpv* loci. Local climate could indeed play a role in the emergence of strains capable of breaking down *Rpv* loci. The Cilaos cirque, situated at an elevation of approximately 600–1,300 masl and surrounded by dense forests, has a humid climate with rainfall throughout the year, more abundant in summer, and cool but not cold winters (Climate Data - France, 2025; du Rau, 1956; Morel et al., 2014). This is characteristic of a temperate oceanic climate according to the Köppen–Geiger classification, which can more broadly be described as subtropical (Climate Data - France, 2025; Trewartha and Horn, 1980). These climatic conditions are highly conducive to grapevine downy mildew epidemics (Santos et al., 2020).

Regarding grapevine cultivation and resistance sources in Cilaos, vineyard areas on Réunion Island have always remained limited, largely due to the challenges posed by the subtropical climate, which hampers dormancy breaking and flower induction (Hermann, 1885). Since the introduction of grapevine downy mildew to the island in the late 19^th^ century, growers have relied on French– American hybrids that naturally exhibit partial resistance to the pathogen, some of which carry the *Rpv3.1* locus. More recently, in 2019–2020, INRAE-ResDur1 varieties combining *Rpv1* and *Rpv3.1* resistance loci were planted, followed shortly afterward, in 2020, by German-bred varieties such as Solaris and Muscaris, carrying *Rpv10* (Torregrosa et al., 2024). As a result, all three loci — *Rpv1*, *Rpv3.1*, and *Rpv10* — are now present in Cilaos, exerting selective pressure on *P. viticola*. Furthermore, between 2019 and 2023, several vineyards in Cilaos were reported to have received no phytosanitary treatments, which likely contributed to the severity of disease outbreaks (O. Yobrégat, personal communication, 2025).

Consistent with its virulent phenotype, the Cilaos strain exhibits a homozygous deletion of effector genes at the *AvrRpv3.1* locus. In addition, it harbors the *avr1* allele, one of the most prevalent virulence alleles in France (Paineau et al., 2024), suggesting that this allele may have been introduced into Réunion Island through a founder population from mainland France, consistent with our population genetic structure analyses. Nonetheless, we cannot completely rule out the possibility of an independent adaptation, with mutations converging into the *avr1* allele, given the long-lasting cultivation of *Rpv3.1*-carrying hybrids in Cilaos.

The breakdown of *Rpv10* in the Cilaos strain presents a different narrative. Several lines of evidence suggest that the *Rpv10* breakdown observed in this study is fundamentally distinct from that described in Europe by Paineau et al. (2022) and Dvorak et al. (2025a). First, the HR triggered by the plant is not completely suppressed in European strains, leading to only partial breakdown of resistance. In contrast, infection with the Cilaos strain produced no necrosis in any repetition. Second, the Cilaos strain lacked the known suppressor allele (S-*AvrRpv10*) previously associated with *Rpv10* breakdown in populations from the Upper Rhine Plain in Europe, that resulted from an admixture event with populations of American origin (Dvorak et al., 2025a). Our analyses indicate that the Cilaos strain has no American genetic background and is genetically distant from both North American populations and the *Rpv10* breaking isolate (Pv1419_1) of German origin (Figure 6, Supplementary Figure 6). Finally, the Cilaos strain harbors a novel mutation at the S-*AvrRpv10* locus described by Dvorak et al. (2025a), featuring a large homozygous deletion. Together, these findings support the hypothesis of a local and independent breakdown of *Rpv10* in Cilaos, likely involving a novel virulence mechanism remaining to be identified.

The breakdown of *Rpv10* by the Cilaos strain was unexpected, as this strain was not collected from a plot containing *Rpv10* varieties, raising questions about its origin. The vineyard of origin is located 4km (as the crow flies) from the nearest vineyard planted with *Rpv10*-carrying varieties (O. Yobrégat, personal communication, 2025). Although *P. viticola* is reported to disperse over several tens of meters (Gobbin et al., 2007, 2005), it may be capable of long-distance aerial dispersal, as observed in other oomycete pathogens at continental scales (Campbell and Ristaino, 1999). An alternative explanation is that the breakdown of *Rpv10* and *Rpv1* in the Cilaos strain represents a case of cross-infectivity. According to the definition of Moury et al. (2014), the mutation conferring virulence on *Rpv1* varieties could also confer virulence on *Rpv10*, with selective pressure from *Rpv1* being responsible for the *Rpv10* breakdown. This could explain why the Cilaos strain, collected from a variety carrying *Rpv1* and *Rpv3.1*, is also virulent on *Rpv10*. Confirming this hypothesis will require identifying the virulence determinants and further studying the *P. viticola* population in Cilaos. The origin of *Rpv10* breakdown in the Cilaos strain, therefore, remains unresolved.

The adaptation of *P. viticola* to host resistance (*Rpv1* and *Rpv10*) occurred on Réunion Island despite the very limited deployment of resistant varieties and within a remarkably short time frame. Assuming independent breakdown events and disregarding the cross-infectivity hypothesis, virulence against *Rpv1* and *Rpv10* emerged only four and three years, respectively, after the introduction of varieties carrying these loci in 2019 and 2020. The fast local adaptation capacity of *P. viticola* has been documented before, including resistance to fungicides (Chen et al., 2007; Delmas et al., 2017) as well as for the breakdown of *Rpv3.1* which occurred in multiple locations within just five years (Delmotte et al., 2014).

In addition to genomic plasticity (Dussert et al., 2019; Dvorak et al., 2025b; Foria et al., 2020), the evolutionary potential of *P. viticola* is further enhanced by large population sizes and a mixed reproductive strategy involving both asexual and sexual reproduction (Gessler and Pertot, 2011). During the grapevine growing season, *Rpv* loci confer partial resistance to *P. viticola* by limiting its growth and spread, but at fall, the pathogen can undergo sexual reproduction on resistant varieties allowing recombination between existing genotypes (Delbac et al., 2019). This genetic exchange increases standing variation in the pathogen population for the following season and can recombine different virulences into one genotype. These dynamics underscore the importance of continuous surveillance of pathogen populations across seasons and, more importantly, highlight the need for new disease management strategies that limit sexual reproduction.

Our findings raise important concerns for the sustainable management of resistant grapevine varieties. They highlight a potential risk of a similar *Rpv1* breakdown in continental Europe and the emergence of more strains able to breakdown multiple *Rpv* loci. This is particularly critical in France where grapevine breeding programs rely on pyramiding *Rpv1* with *Rpv3.1* and *Rpv10* (program ResDur3)(Avia et al., 2023). While the less conducive climatic conditions in Europe and the recommended targeted phytosanitary treatments on resistant varieties may reduce the risk of multivirulent strain emergence, evidence from 2024 prompts vigilance. An increase in disease incidence was observed in vineyards planted with the Artaban variety (Rpv1–Rpv3.1) in southeastern France, and subsequent phenotyping assays confirmed that several strains were capable of overcoming *Rpv1*-mediated resistance. (Pelissier et al., 2025). These events, however, remain localized, with outbreaks detected only in that region (Observatoire national du déploiment des cépages résistants, 2025). Overall, vineyards planted with *Rpv1*-carrying varieties, such as Bouquet and ResDur lines, continue to benefit from effective resistance conferred by this locus.

Nevertheless, these observations underscore the need for ongoing monitoring and further investigation of *Rpv1* breakdown.

The emergence of multi-virulent *P. viticola* strains capable of overcoming major resistance loci including *Rpv1*, *Rpv3.1*, and *Rpv10*, raises serious concerns about the long-term durability of a strategy relying solely on resistance loci pyramiding. This highlights that sustainable management of resistant grapevine varieties must combine intensive field monitoring of grapevine downy mildew epidemics with laboratory-based phenotyping to detect emerging adaptations. Achieving this requires coordinated monitoring networks, standardized methods, reference strains for phenotyping, and stakeholders alert protocols. When integrated with population genomics, such effort can identify *avr* loci and the genetic mechanisms underlying plant–pathogen interactions, improving our understanding of evolutionary dynamics and enabling early detection of virulence. This underscores the need for future research to pinpoint the genetic determinants of *Rpv1* and *Rpv10* breakdown on Réunion Island, using approaches such as GWAS or QTL mapping, which have proven effective in this pathosystem (Dvorak et al., 2025a; Paineau et al., 2024). Without such coordinated efforts, decades of breeding could risk being rendered ineffective.

## Conclusion

We report the breakdown of *Rpv1* by *P. viticola* in Cilaos, characterized the first strain capable of overcoming three *Rpv* loci, and provided evidence for a novel mechanism underlying the breakdown of *Rpv10*. Our findings raise important concerns for viticulture in Europe. The emergence of this multi-virulent strain that shares genetic traits with European populations, suggests that similar breakdowns could soon occur on Europe. While this represents a significant threat to grapevine disease management, our study also provides a timely opportunity to anticipate and respond to such host-adaptation of the pathogen. Beyond its practical implications for resistance monitoring and breeding strategies, this work makes a significant contribution to our understanding of host-pathogen interactions and the evolutionary dynamics that shape the durability of plant disease resistance. Furthermore, it lays the groundwork for further investigation on the identification of the virulence determinants of these new resistance breakdowns.

## Declarations

### Availability of data and materials

The sequencing data of the Cilaos strain produced in this study has been deposited under BioProject PRJNA1255592.

### Competing interests

The authors declare that they have no competing interests.

### Funding

This study was funded by the Plant Health and Environment Division of the French National Research Institute for Agriculture, Food and Environment (INRAE), the French National Research Agency (PPR VITAE, grant 20-PCPA-0010), and the French government as part of France 2030 (Project BPI France, Innovitiplant).

### Authors’ contributions

J.R.M.: Drafting of the manuscript; analysis of phenotypic and genomic data.

A.-S.M.: Lead of phenotypic experiments; analysis of phenotypic results; critical review of the manuscript.

E.D.: Population structure analysis of the largest strain dataset; critical review of the manuscript; photos and guidance for the description of visual scale of necrosis.

I.D.M.: Carried out experiments and provided technical expertise. C.C.: Management of genomic data.

L.D.: Lead of phenotypic experiments. F.F.: Critical review of the manuscript.

I.H.: Field monitoring and isolation of the strain from Cilaos, Réunion Island.

O.Y.: Field monitoring and isolation of the strain from Cilaos, Réunion Island; contribution to drafting of the discussion.

M.F.-O.: Project leadership; drafting and critical review of the manuscript. F.D.: Project leadership; drafting and critical review of the manuscript.

## Acknowledgements

We thank the Genotoul Bioinformatics Platform in Toulouse, France, for supplying the computing and storage resources, and to the staff in our research unit (SAVE) for their help during experiments.

## Supplementary material

### Supplementary tables

**Supplementary table 1.** *Plasmopara viticola* strains included in a population structure analysis, t<u>heir geographical origin and accession to genomic sequencing data.</u>.

| ID Strain | Country | Location | Genomic sequencing data |
| --- | --- | --- | --- |
| Pv8586 | France | Cilaos – Réunion Island | PRJNA1255592 |
| Pv221_1 | France | Blanquefort | PRJNA329579 |
| Pv2128_1 | France | Villenave d'Ornon | PRJNA1255592 |
| Pv2254_1 | France | Chitry | PRJNA1255592 |
| Pv2578_1 | France | Le Landreau | PRJNA1255592 |
| Pv2664_1 | France | Villenave d'Ornon | PRJNA1255592 |
| Pv2868_1 | France | Pontèves | PRJNA1255592 |
| Pv3112_1 | France | Gruissan | PRJNA1255592 |
| Pv3334 | France | Rodhilar | PRJNA1255592 |
| Pv3337 | France | Rodhilar | PRJNA1255592 |
| Pv4089_1 | France | Villenave d'Ornon | PRJNA875296 |
| Pv4160_1 | France | Villenave d'Ornon | PRJNA875296 |
| Pv4742 | France | Romagne | PRJNA1255592 |
| Pv7219_1 | France | Wintzenheim | PRJNA1255592 |
| Pv8786_1 | France | Bieujac | PRJNA1255592 |
| Pv2219_1 | Spain | Sant Sadurni D'Anoia | PRJNA1255592 |
| Pv2221_1 | Spain | Sant Sadurni D'Anoia | PRJNA1255592 |
| Pv2596_11 | Spain | Agoncillo | PRJNA1255592 |
| Pv412_11 | Switzerland | Cugnasco | PRJNA1095879 |
| Pv413_11 | Switzerland | Cugnasco | PRJNA644583 |
| VB = Pv8522_1 | Switzerland | Soyhières | PRJNA875296 |
| Pv8592 | Switzerland | Soyhières | PRJNA1255592 |
| Pv8593 | Switzerland | Soyhières | PRJNA1255592 |
| Pv1174_1 | Germany | Ihringen | PRJNA644583 |
| Pv1238_1 | Germany | Pfaffenweiler | PRJNA644583 |
| Pv1286_1 | Germany | Ehrenkirchen | PRJNA644583 |
| Pv1314_1 | Germany | Ebringen | PRJNA644583 |
| Pv1419_1 | Germany | Ihringen | PRJNA1095879 |
| Pv9239 | Germany | Freiburg | PRJNA1255592 |
| Pv9949 | Germany | Eichstetten | PRJNA1255592 |
| Pv336 | Hungary | Eger | PRJNA644583 |
| Pv340_1 | Hungary | Tolcsva | PRJNA644583 |
| Pv2543_1 | Hungary | Pécs | PRJNA1095879 |
| Pv2546_1 | Hungary | Pécs | PRJNA1255592 |
| Pv2547_1 | Hungary | Pécs | PRJNA1255592 |
| Pv2924 | Hungary | Pécs | PRJNA1255592 |
| Pv2938 | Hungary | Pécs | PRJNA1255592 |
| Pv2955 | Hungary | Pécs | PRJNA1255592 |
| Pv2958 | Hungary | Pécs | PRJNA1255592 |
| Pv2959 | Hungary | Pécs | PRJNA1255592 |
| Pv2961 | Hungary | Pécs | PRJNA1255592 |
| Pv2963 | Hungary | Pécs | PRJNA1255592 |
| Pv2966 | Hungary | Pécs | PRJNA1255592 |
| Pv2967 | Hungary | Pécs | PRJNA1255592 |
| Pv2972 | Hungary | Pécs | PRJNA1255592 |
| Pv2416_1 | Italy | Pianello Val Tidone | PRJNA644583 |
| Pv2821_1 | Italy | Piacenza | PRJNA1255592 |
| Pv3174_11 | Italy | Piateda | PRJNA644583 |
| Pv3175_1 | Italy | Canevino | PRJNA644583 |
| Pv9113_1 | Italy | North-East | PRJNA1255592 |
| Pv9173_1 | Italy | North-East | PRJNA1255592 |
| Pv9189_1 | Italy | North-East | PRJNA1255592 |
| Pv2534_1 | Lebanon | NC | PRJNA1255592 |
| Pv3190_1 | Georgia | NC | PRJNA644583 |
| YL | China | Yangling, Shaanxi province | PRJNA488341 |
| PvCAN1 | Canada | Quebec province | PRJNA1255592 |
| PvCAN2 | Canada | Quebec province | PRJNA1255592 |
| USDA-Pv3 | United States | Geneva, NY | PRJNA1255592 |

**Supplementary table 2.** Results of a pathogenicity assay in which nine *Plasmopara viticola* strains were inoculated into leaf discs of different grapevine varieties: Chardonnay (Susceptible), 3179-90-7 (*Rpv1*), Regent (*Rpv3.1*), Muscaris (*Rpv10*) and Fleurtai (*Rpv12*). Values of mean sporulation area, mean Hypersensitive Response (HR) and mean Efficiency of the Resistance Factor (ERF) index are displayed for each combination strain-variety.

| Variety | Strain | mean sporulation<br>area (%) | mean HR | mean ERF<br>index |
| --- | --- | --- | --- | --- |
| Chardonnay | Pv8586_1 | 4.263 | 0.000 | NA |
| Chardonnay | Pv8522_1 | 9.465 | 0.833 | NA |
| Chardonnay | Pv412_11 | 15.160 | 0.167 | NA |
| Chardonnay | Pv3174_11 | 7.104 | 0.167 | NA |
| Chardonnay | Pv2909_11 | 16.222 | 0.000 | NA |
| Chardonnay | Pv2664_1 | 14.339 | 0.667 | NA |
| Chardonnay | Pv2578_1 | 10.410 | 0.000 | NA |
| Chardonnay | Pv2543_1 | 16.679 | 0.500 | NA |
| Chardonnay | Pv1419_1 | 15.670 | 1.000 | NA |
| 3179-90-7 | Pv8586_1 | 7.156 | 0.000 | -0.679 |
| 3179-90-7 | Pv8522_1 | 0.518 | 3.000 | 0.945 |
| 3179-90-7 | Pv412_11 | 8.611 | 2.000 | 0.432 |
| 3179-90-7 | Pv3174_11 | 1.955 | 2.833 | 0.725 |
| 3179-90-7 | Pv2909_11 | 3.177 | 2.833 | 0.804 |
| 3179-90-7 | Pv2664_1 | 4.294 | 2.500 | 0.701 |
| 3179-90-7 | Pv2578_1 | 3.526 | 3.000 | 0.661 |
| 3179-90-7 | Pv2543_1 | 3.759 | 2.000 | 0.775 |
| 3179-90-7 | Pv1419_1 | 8.056 | 2.000 | 0.486 |
| Fleurtaï | Pv8586_1 | 0.000 | 5.000 | 1.000 |
| Fleurtaï | Pv8522_1 | 18.745 | 1.000 | -0.981 |
| Fleurtaï | Pv412_11 | 0.031 | 3.000 | 0.998 |
| Fleurtaï | Pv3174_11 | 0.000 | 4.500 | 1.000 |
| Fleurtaï | Pv2909_11 | 0.000 | 4.667 | 1.000 |
| Fleurtaï | Pv2664_1 | 0.000 | 3.167 | 1.000 |
| Fleurtaï | Pv2578_1 | 0.000 | 4.000 | 1.000 |
| Fleurtaï | Pv2543_1 | 16.657 | 0.667 | 0.001 |
| Fleurtaï | Pv1419_1 | 0.001 | 3.000 | 1.000 |
| Muscaris | Pv8586_1 | 3.247 | 0.000 | 0.238 |
| Muscaris | Pv8522_1 | 0.000 | 4.000 | 1.000 |
| Muscaris | Pv412_11 | 0.005 | 3.000 | 1.000 |
| Muscaris | Pv3174_11 | 0.000 | 4.500 | 1.000 |
| Muscaris | Pv2909_11 | 0.000 | 4.667 | 1.000 |
| Muscaris | Pv2664_1 | 0.000 | 3.500 | 1.000 |
| Muscaris | Pv2578_1 | 0.000 | 3.667 | 1.000 |
| Muscaris | Pv2543_1 | 0.000 | 3.500 | 1.000 |
| Muscaris | Pv1419_1 | 12.632 | 1.333 | 0.194 |
| Regent | Pv8586_1 | 12.886 | 0.000 | -2.023 |
| Regent | Pv8522_1 | 17.592 | 0.667 | -0.859 |
| Regent | Pv412_11 | 33.089 | 0.167 | -1.183 |
| Regent | Pv3174_11 | 2.319 | 3.000 | 0.674 |
| Regent | Pv2909_11 | 2.514 | 3.000 | 0.845 |
| Regent | Pv2664_1 | 19.686 | 1.000 | -0.373 |
| Regent | Pv2578_1 | 2.951 | 3.000 | 0.717 |
| Regent | Pv2543_1 | 9.633 | 2.167 | 0.422 |
| Regent | Pv1419_1 | 19.343 | 1.000 | -0.234 |

**Supplementary table 3.** Results of a pathogenicity assay in which nine *Plasmopara viticola* strains were inoculated into leaf discs of the 5^th^, 8^th^ and 10^th^ below the apex of different grapevine varieties: Chardonnay (Susceptible), 3179-90-7 (*Rpv1*), and Regent (*Rpv3.1*). Values of mean sporulation area, mean Hypersensitive Response (HR) and mean Efficiency of the Resistance <u>Factor (ERF) index are displayed for each combination strain-variety.</u>.

| Variety | Strain | mean sporulation area (%) | mean HR | mean ERF Index |
| --- | --- | --- | --- | --- |
| Chardonnay | Pv8586_1 | 22.121 | 0.000 | NA |
| Chardonnay | Pv8522_1 | 20.466 | 0.667 | NA |
| Chardonnay | Pv3174_11 | 2.652 | 0.000 | NA |
| Chardonnay | Pv2909_11 | 21.304 | 0.000 | NA |
| Chardonnay | Pv2664_1 | 25.569 | 1.500 | NA |
| Chardonnay | Pv2578_1 | 17.588 | 0.000 | NA |
| Chardonnay | Pv2543_1 | 13.927 | 0.333 | NA |
| Chardonnay | Pv1419_1 | 20.839 | 0.500 | NA |
| 3179-90-7 | Pv8586_1 | 33.233 | 0.000 | -0.550 |
| 3179-90-7 | Pv8522_1 | 2.594 | 3.000 | 0.903 |
| 3179-90-7 | Pv3174_11 | 1.165 | 3.167 | 0.560 |
| 3179-90-7 | Pv2909_11 | 2.756 | 3.333 | 0.826 |
| 3179-90-7 | Pv2664_1 | 1.276 | 4.000 | 0.862 |
| 3179-90-7 | Pv2578_1 | 0.782 | 3.500 | 0.960 |
| 3179-90-7 | Pv2543_1 | 3.171 | 2.667 | 0.799 |
| 3179-90-7 | Pv1419_1 | 6.619 | 3.000 | 0.674 |
| Regent | Pv8586_1 | 27.272 | 0.000 | -0.166 |
| Regent | Pv8522_1 | 29.630 | 1.333 | -0.138 |
| Regent | Pv3174_11 | 0.241 | 4.500 | 0.937 |
| Regent | Pv2909_11 | 3.150 | 3.667 | 0.916 |
| Regent | Pv2664_1 | 20.337 | 0.833 | 0.148 |
| Regent | Pv2578_1 | 0.868 | 4.000 | 0.973 |
| Regent | Pv2543_1 | 2.425 | 3.667 | 0.911 |
| Regent | Pv1419_1 | 23.892 | 0.500 | 0.322 |

## Supplementary figures

**Supplementary Figure 1.**
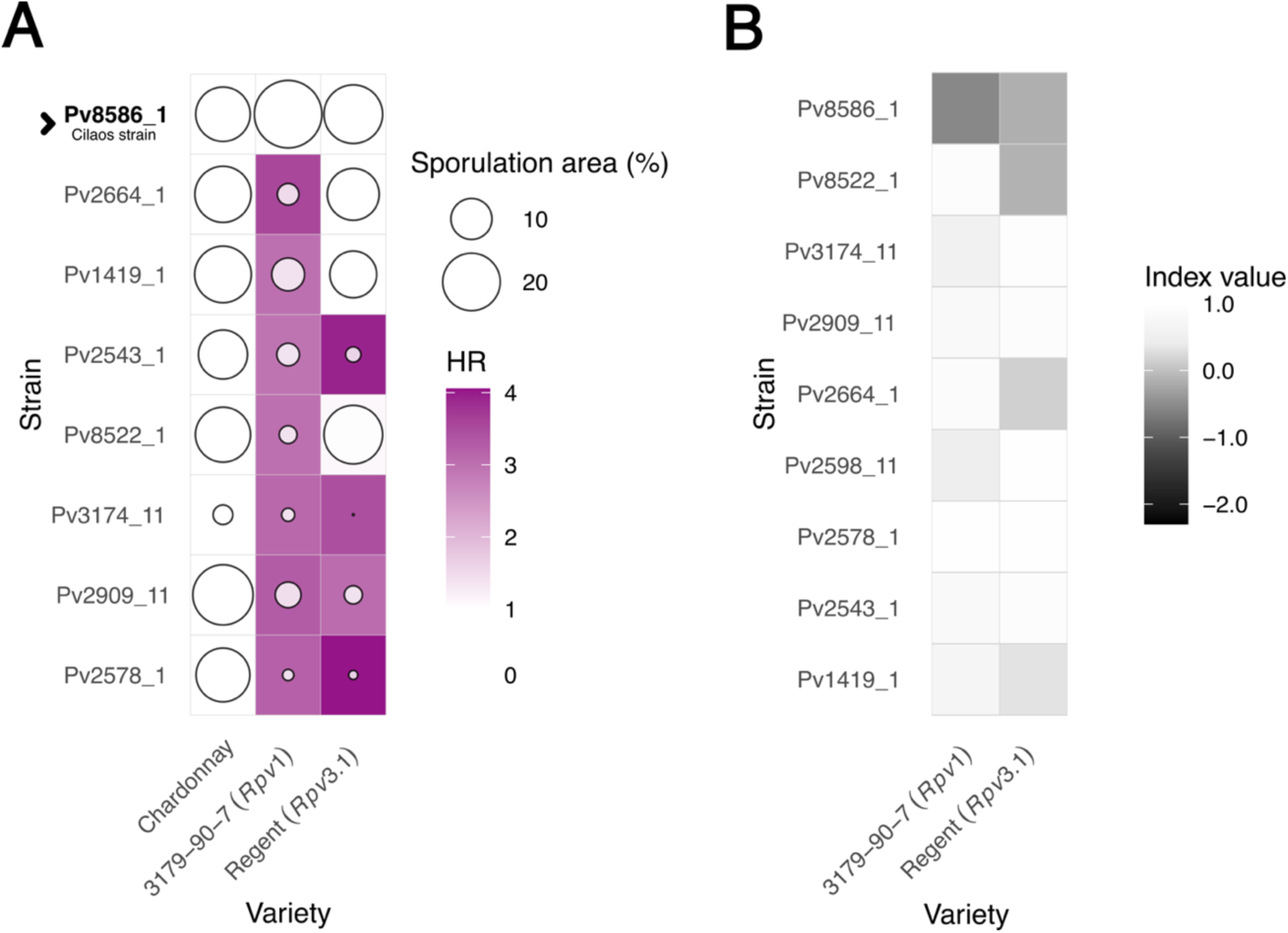
Results of a second cross-inoculation experiment showing a *Plasmopara viticola* strain from Cilaos – Réunion Island able to breakdown the resistance conferred by the *Rpv1 locus* on grapevine, validated by the use of a reference pool of strains of the same species. All strains were inoculated on varieties carrying either the resistance factors *Rpv1* or *Rpv3.*1 and on a susceptible variety (Chardonnay). Values of HR and sporulation were determined after the incubation period post-inoculation in different leaf stages, 5^th^, 8^th^ and 10^th^ then values were averaged. **A.** The strain names are displayed alongside a heatmap presents the percentage of sporulating area and the Hypersensitive Response (HR) score for each interaction between Variety and Strain. For the HR, the heatmap shows the mean value of the repetitions, represented on a color scale. For the sporulating area, values are represented by circles, with the size of each circle corresponding to the percentage value. **B.** Heatmap that shows the values of Efficiency of the Resistance Factor (ERF index) for each interaction tested, except for the susceptible variety.

**Supplementary figure 2.**
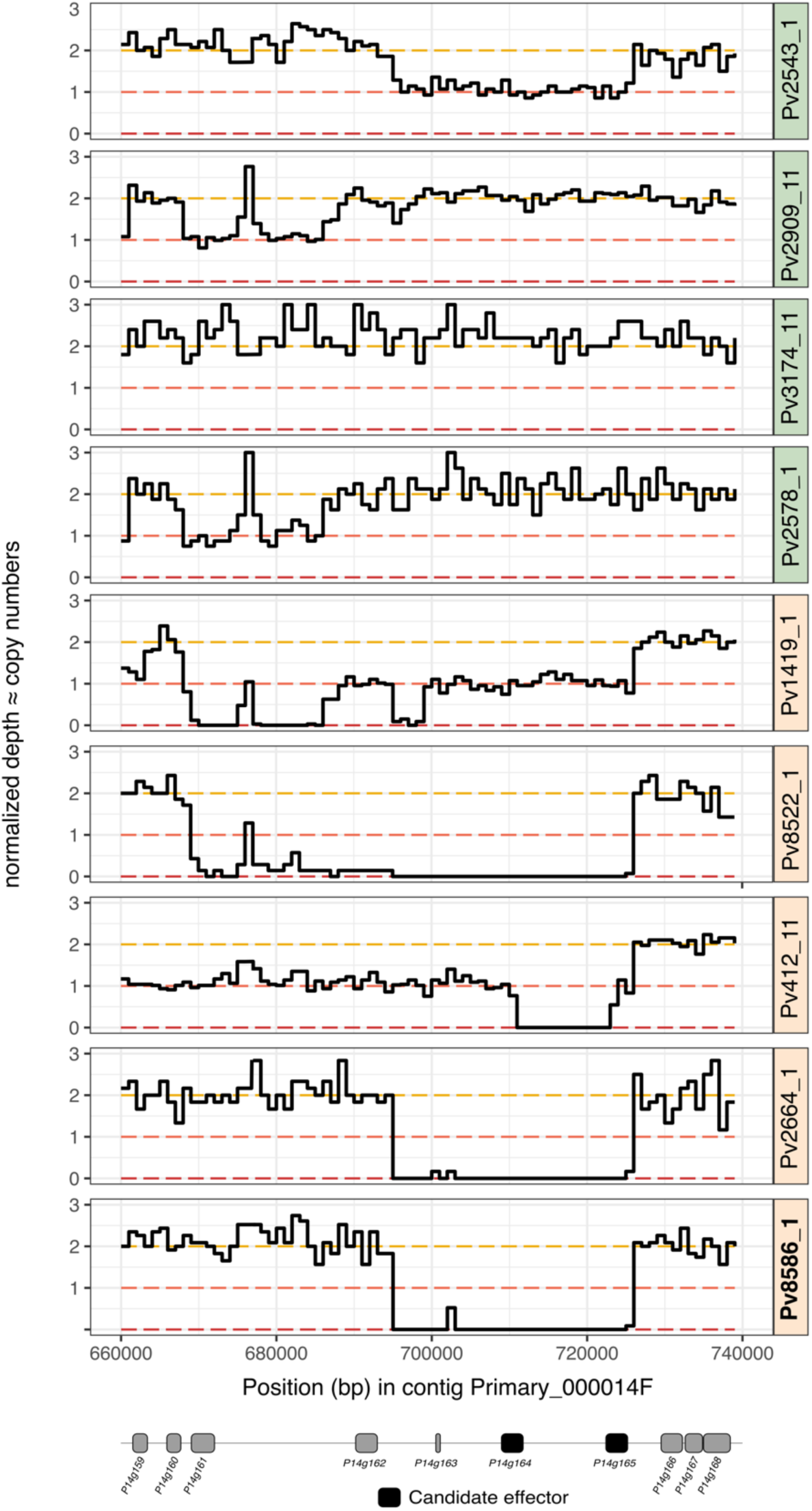
Read depth of sequencing at the *AvrRpv3.1* locus in avirulent and virulent *P. viticola* strains. Normalized read depth is displayed along the genomic region. Estimated copy number is indicated on the y-axis and calculated by 1kb windows. At the bottom, genes are indicated as boxes, where deep-magenta boxes indicate candidate effectors. Avirulent strains names are enclosed on a green box whereas virulent strains are enclosed in a light-peach colored box. Strain coming from Cilaos on Réunion Island is written in bold letters.

**Supplementary figure 3.**
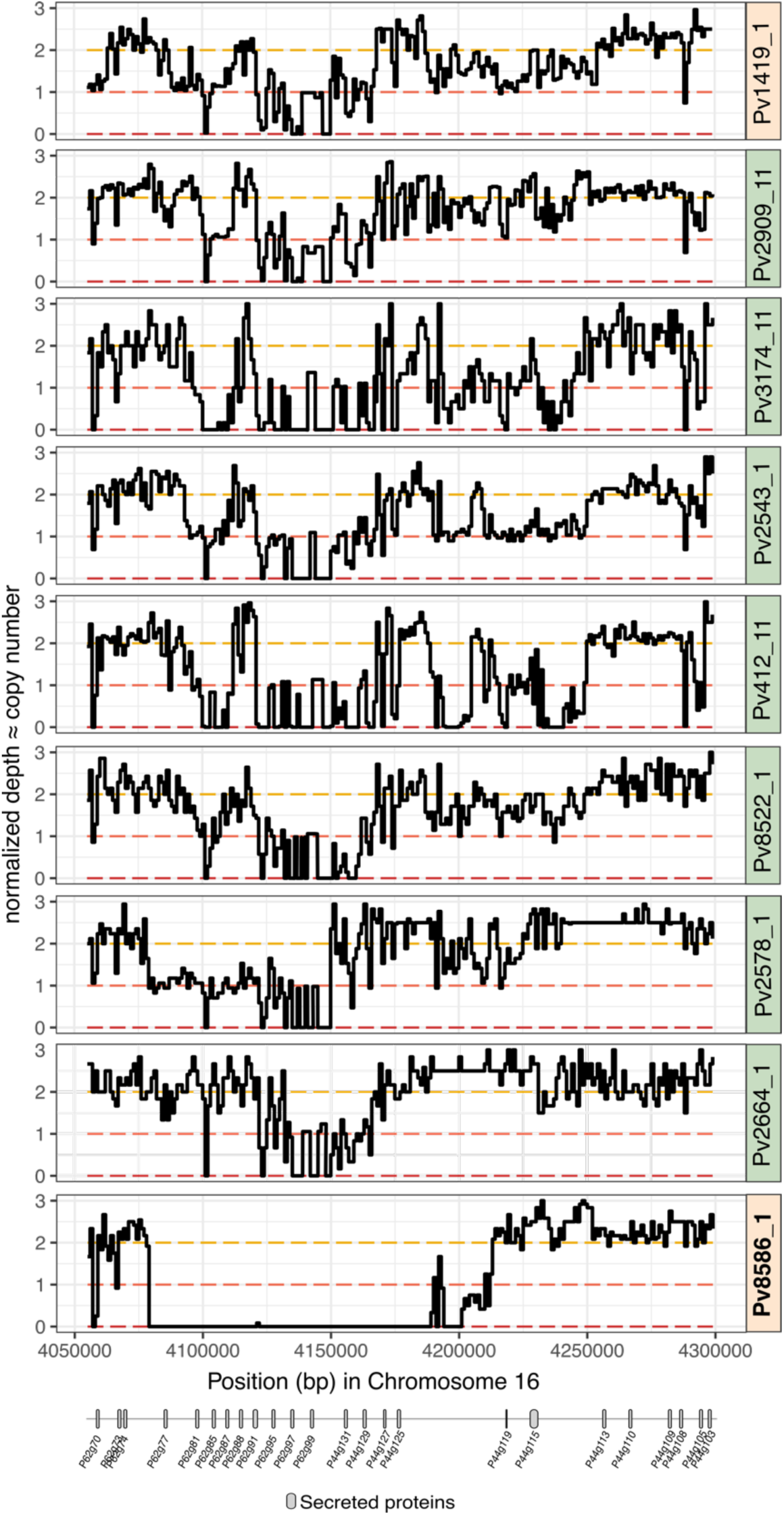
Read depth of sequencing at the S-*AvrRpv10* locus in avirulent and virulent *P. viticola* strains when mapped to the Haplotype 1 of the virulent reference strain Pv1419_1. Normalized read depth is displayed along the genomic region. Estimated copy number is indicated on the y-axis and calculated by 4kb windows. At the bottom, genes are indicated as boxes, where deep-magenta boxes indicate candidate effectors. Avirulent strains names are enclosed on a green box whereas virulent strains are enclosed in a light-peach colored box. Strain coming from Cilaos on Réunion Island is written in bold letters. All strains, except the Cilaos strain, show coverage across the entire studied genomic region.

**Supplementary figure 4.**
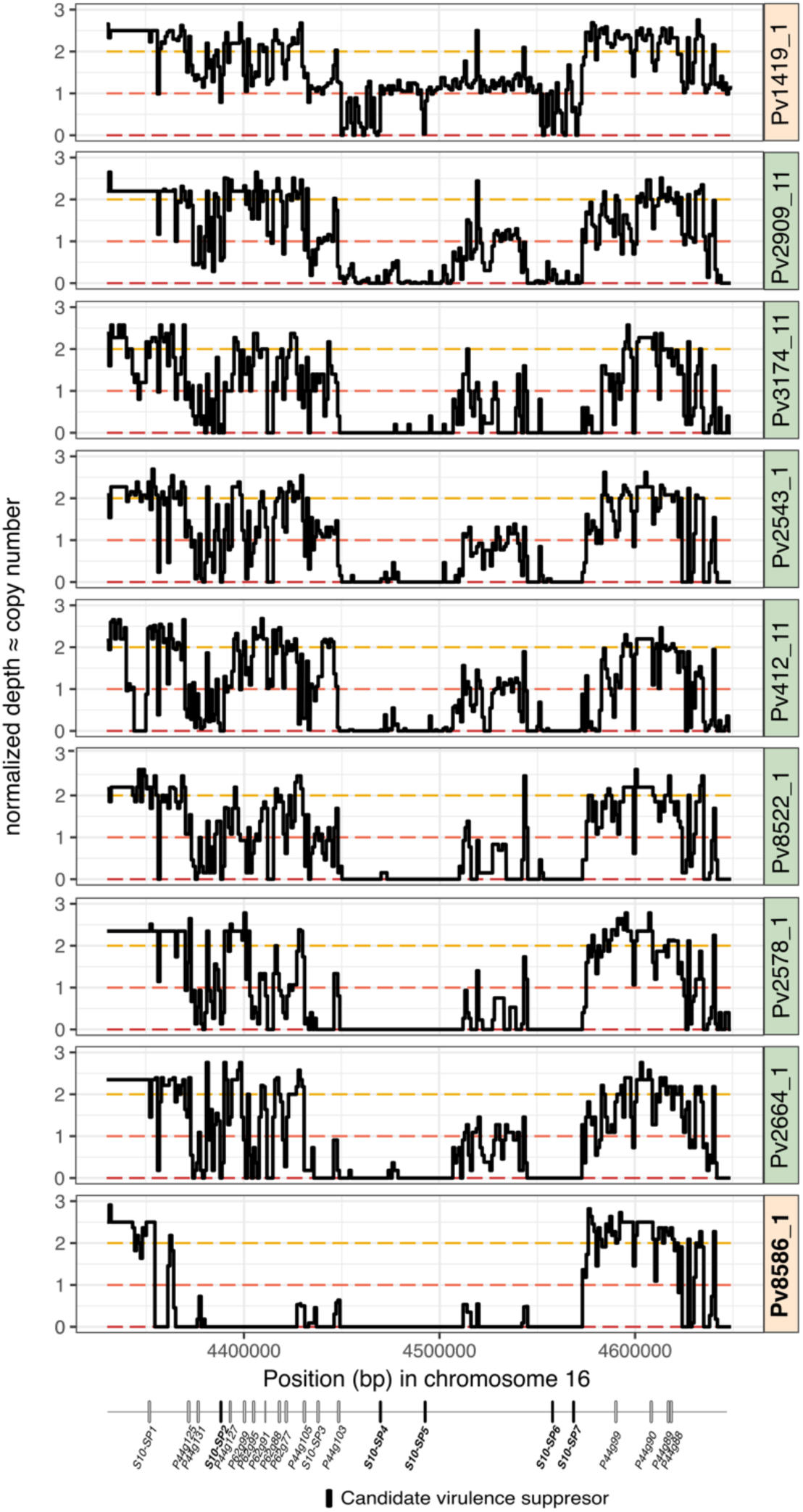
Read depth of sequencing at the S-*AvrRpv10* locus in avirulent and virulent *P. viticola* strains when mapped to the Haplotype 2 of the virulent reference strain Pv1419_1. Normalized read depth is displayed along genomic region. Estimated copy number is indicated on the y-axis and calculated by 4kb windows. At the bottom, genes are indicated as boxes, where deep-magenta boxes indicate candidate effectors. Avirulent strains names are enclosed on a green box whereas virulent strains are enclosed in a light-peach colored box. Strain coming from Cilaos on Réunion Island is written in bold letters. Only the strain Pv1419_ show coverage across the entire studied genomic region, the rest of the strains present only some genes, and the Cilaos strain does not harbor most of the studied genomic region.

**Supplementary figure 5.**
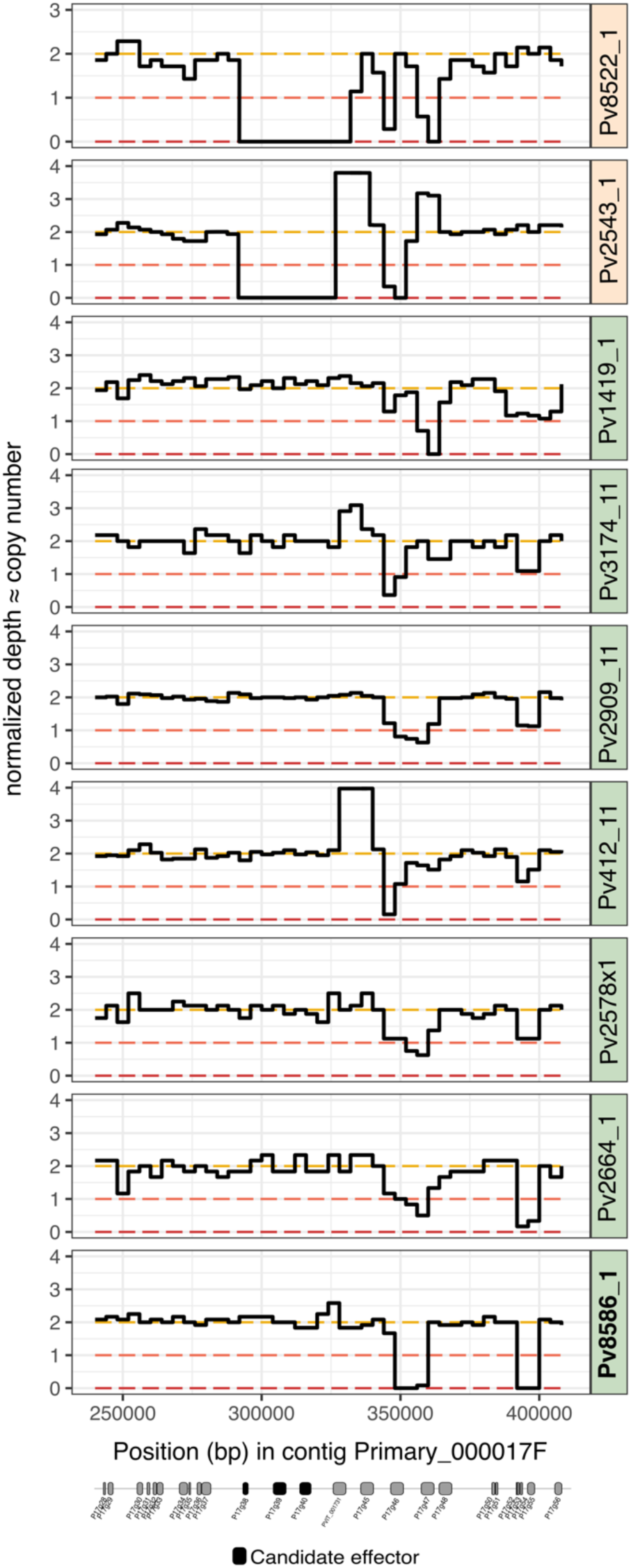
Read depth of sequencing at the *AvrRpv12* locus in avirulent and virulent *P. viticola* strains. Normalized read depth is displayed along the genomic region. Estimated copy number is indicated on the y-axis and calculated by 1kb windows. Some strains present higher coverage with more than two copies, likely due to additional copies of genes. At the bottom, genes are indicated as boxes, where deep-magenta boxes indicate candidate effectors. Avirulent strains names are enclosed on a green box whereas virulent strains are enclosed in a light-peach colored box. Strain coming from Cilaos on Réunion Island is written in bold letters.

**Supplementary figure 6.**
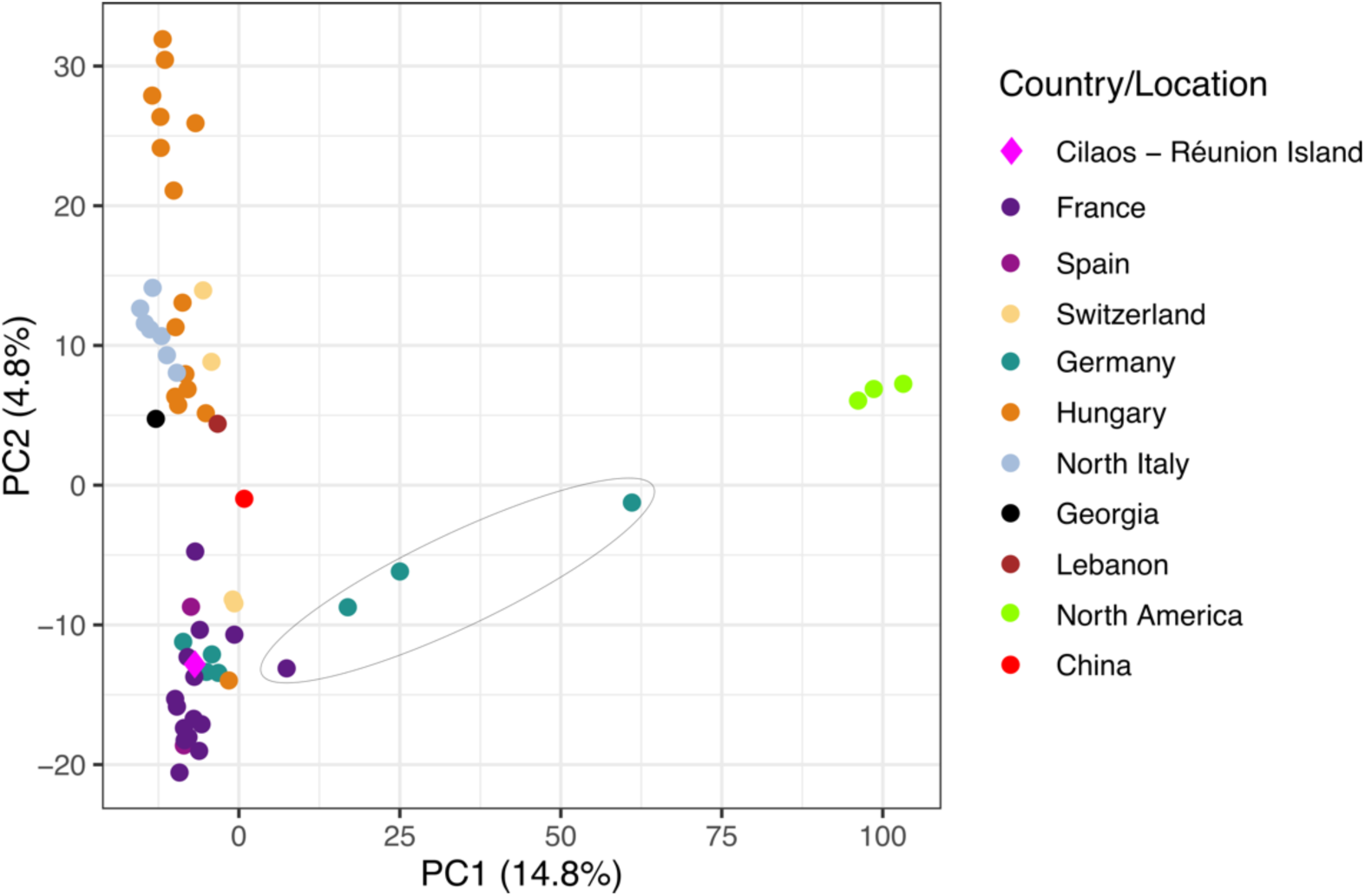
Principal Component Analysis (PCA) of 89,673 SNPs from 58 *Plasmopara viticola* strains, showing that a strain from Cilaos (Réunion Island) clusters closely with strains from continental Europe, particularly those from the western side of the Alps (France, Spain, and Germany). Closest strains are French. The SNPs were obtained using the same dataset and methodology as described in Dvorak et al. (2025a https://doi.org/10.1101/2025.05.18.654733), but adding the genomic information of the Cilaos strain. Analysis included the strain from Cilaos plus strains from Canda (n=2), United States of America (n=1), China (n=1), Georgia (n=1), Lebanon (n=1), France(n=15), Hungary (n=15), Switzerland (n=5), Italy (n=7) and Germany (n=7). Ellipse encloses strains that are admixed with North-American populations and are virulent on *Rpv10*-carrying varieties.

## Notes

### Competing Interest Statement

The authors have declared no competing interest.

